# Kaolinite deposits in the upper Iguaçu river, Brazil: formation and mineralogical attributes

**DOI:** 10.1101/2021.03.04.433860

**Authors:** Daniela N. Ferreira, Vander de Freitas Melo, Pablo Vidal-Torrado, Jairo Calderari Oliveira

## Abstract

Kaolinite (Kt) is the most studied mineral, being widely used in the ceramic, pharmaceutical and cellulose industries, in addition to being the main soil mineral in the world. Found in different parts of the planet, it differs in genesis and may be formed as a result of local weathering of the rocks, occurring in the silt fraction; and also due to the mineral’s neogenesis with a predominance of clay-sized particles. The plain of upper Iguaçu river has the largest kaolinitic deposit in the south of Brazil and it’s formation raised doubts if this kaolin was transported or formed in situ due the high organic matter in the alluvial plain. To elucidate the origin of kaolin deposits, we sampled a possible font of the mineral, in the mountains of Serra do Mar and sampled two tubes that reach 4 m depth in the upper Iguaçu plain. We performed textural analysis, organic carbon, X-ray diffraction, Kt crystallography in silt and clay fractions, thermal analysis (TDA/TG) to quantify Kt and Gb in the clay fraction and scanning electron microscopy (SEM) with dispersive energy spectroscopy (EDS). The TDA/TG analysis demonstrated that saprolite has 66% of the kaolinite found in the plain. The XRD analysis shows a significant presence of mica (Mc) in all samples of the silt fraction, both in Serra do Mar and in the plain. In SEM/EDS, crystals with planar growth are observed, and the presence of pseudomorphic Mc-Kt in the silt fraction of all analyzed samples, with emphasis on the tubes sample with the crystal having almost twice the size of that observed in the saprolite sample from Serra do Mar, allowing to infer that the Kt of the silt fraction of the wetland soils were formed on site by the diagenesis of mica particles. The results obtained in this work indicate that the kaolinitic material found in the wetland of the upper Iguaçu plain is the result of weathering processes in the wetland itself, evidenced by the large pseudomorphs found, even greater than those observed in Serra do Mar.

## Introduction

Originating from the Chinese word “Kauling”, which means high hill, referring to the hill where it originated, in Jauchau Fu in northern China, kaolin is considered one of the six most abundant minerals in the earth [1, 2]. Kaolin deposits are widely exploited due to their physical and chemical properties, such as manufacturing ceramics, paper, cosmetics, plastics, rubbers, paints, pesticides, food and pharmaceutical products, fertilizers, detergents, cements, among others [3, 4].

Kaolin deposits are found in several areas of the planet, such as the United States [5, 6], Turkey [7], Pakistan [8], Africa [9], Germany [10], Asia [1], South America [11, 12, 13]. Considering the occurrence and its genesis, kaolin can be classified as primary or secondary [9]. Primary kaolin results from the in situ alteration process, either by hydrothermal actions or residual accumulation of the weathering process [1]. In this case it is common the occurrence of the mineral in the silt fraction, mainly by topotaxial alterations of the micas (muscovite), resulting in sericite, which has its edges altered in kaolinite.

Thus, the kaolinite assumes in its genesis the stacking of crystals, characteristic of the group of phyllosilicates, whose growth is guided by microcrystals with a plate format and hexagonal profile [1]. This process is common of rocks such as, for example, granites and rhyolites, which are rich in muscovite, feldspar and Al, where much of the K is removed from the mineral structure, avoiding the formation of illite [1, 14].

On the other hand, secondary kaolin is mostly associated with transport processes and deposition of clasts, of reduced size, predominantly clay-sized particles, deposited in the form of lenses. Another route of secondary kaolin formation is by the weathering of arkose, a coarse material with predominantly feldspathic mineralogy [9].

Brazil has one of the largest kaolinitic clay deposits in the world [12], with significant production in the North region. These deposits were dated as being formed in the Pliocene by Murray [15], while Kotschoubey [16] associate these deposits with events that occurred in the upper Tertiary and Quaternary. Montes et al. [12] and Wilson et al. [13] define these deposits as sedimentary, with strata of varied granulometric composition, indicating various deposition events of kaolinitic material. It is also common to find in these strata, coarser material composed of large amounts of feldspar (arkose), and its weathering could also result in the formation of kaolin.

Through chemical, crystallographic and morphological attributes it is possible to relate minerals from different locations had their genesis in similar conditions [14], establishing the genetic relationship between them and, from this information, work other geological/geomorphological processes.

The plain of upper Iguaçu river is responsible for the largest kaolinitic deposit in the south of the country, estimated at 200 Mt [17]. The geology of the region is formed by rocks of the crystalline basement, more specifically of the granite-gnaisse-migmatite complex. These rocks are found mainly in the part of the relief with the most movement, also called Serra do Mar. In this geomorphological feature it is remarkable the occurrence of orographic rains, which can exceed 2500 mm of annual precipitation, promoting intense alteration and desilication of the rocks mentioned above [18]. The geological and climatic conditions mentioned above are favorable for the formation of kaolinitic deposits in the region.

However, Biondi and Santos [17] postulate the formation of these deposits during the Quaternary, under strong influence of the desilication of deposits of the Guabirotuba Formation. According to these authors, the presence of high levels of organic matter in floodplain areas with a predominance of Histosols would create an acid environment that would increase Si removal and result in kaolinite formation.

On the other hand, Mucha [19] in the same area, found kaolin deposits in well-drained environments, with an abrupt transition to a reddish layer, with higher contents of iron oxides. Still according to this author, these characteristics are typical of sediment transport and deposition processes, raising doubts about the main formation process of kaolinitic deposits and how they could be associated with landscape modeling processes.

Another way of kaolinite formation, without much influence of organic matter, refers to the formation of pseudomorphs, in which biotite or muscovite grains, from the interior to the edge, progressively pass to kaolinite, maintaining the structural continuity [20, 21]. According to these authors, the pseudomorphs of this process can form macrocrystals, with diameter greater than 2 mm, but that with the advancement of the pedogenetic process these crystals would be reduced to less than 2 mm in diameter.

During the transformation process from the mineral present in the rock to the one found in the saprolite or soil, some crystallographic, chemical, or morphological attributes can indicate the intensity of the process. Besides visualizing the formation of pseudomorphs by scanning electron microscopy (SEM), the transformation of 2:1 mineral into kaolinite is identified in X-ray diffractions, using the asymmetry index (AI) of the peak in the plane reflection (001) of kaolinite [22]. Melo et al. [23] observed a strong and direct correlation between AI values with K and Mg contents, suggesting the transformation of biotite into kaolinite.

Also, as a function of the weathering process, the higher iron activity in the solution of the pedogenesis environment can generate isomorphic substitution of Al^3+^ by Fe^3+^, reducing crystallinity and increasing structural disorder and surface area [22]. On the other hand, kaolinite ordering can be studied by some crystallinity indices, such as the Hinckley’s [5] and Hughes and Brown’s [24] crystallinity indices (HBCI). The former has as a major limitation its use in samples with a high degree of disorder, since it is difficult to characterize the peaks of the 02l and 11l planes. The HBCI has as limitation the interference of other minerals that overlap the common peaks of kaolinite [22, 23].

The central hypothesis of this work is that the formation of the kaolinitic deposits in the plain of upper Iguaçu river occurred by deposition of material from the Serra do Mar in a previous geological event, possibly in the Quaternary, with a different environment from the present one. Therefore, the main objective is to compare mineral samples from the two environments, and through qualitative and quantitative analyses of kaolinite, to identify the patterns that reiterate in these environments, in order to corroborate the hypothesis of this work.

## Materials and Methods

### Study area

The study area is located in the metropolitan region of Curitiba, South of Brazil. Geomorphologically it is situated between the Serra do Mar uplifted blocks and the First Paraná Plateau uplifted blocks, both composed of rocks of the gneiss-migmatite complex and, as a result of the pediplanation process, plain of the upper Iguaçu river (Fig 1). The Iguaçu river is of great environmental and economic importance, sheltering extensive floodplain areas in its sources, six hydroelectric plants along its course that supply energy to three countries - Argentina, Paraguay and Brazil.

**Figure 1.**
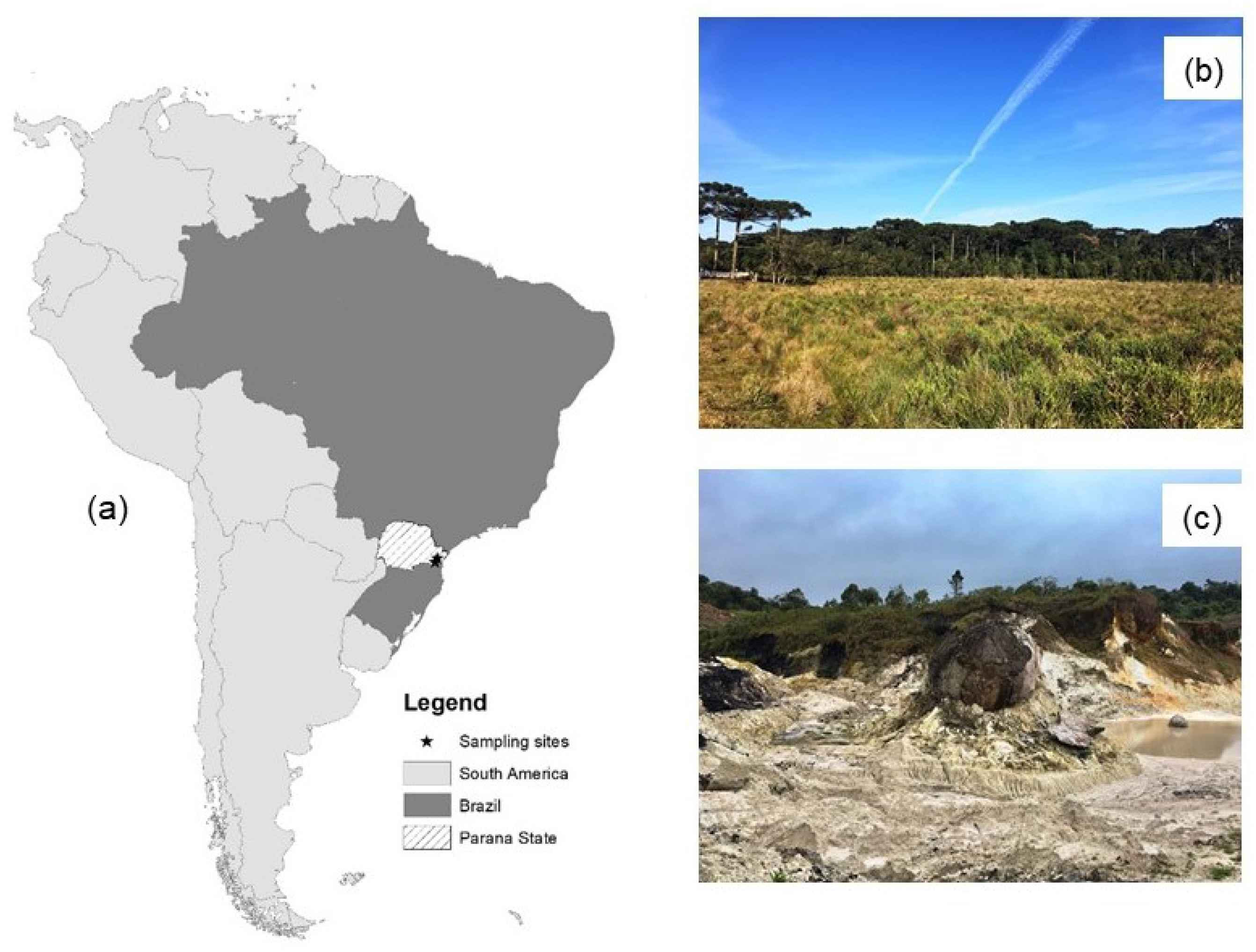
location of the study area and photos of the sampled points. Situation of study site in the South America (a). Plain of the Upper Iguaçu river (b). Kaolin minning in the Serra do Mar (c).

According to Biondi and Santos [17], the plain of the upper Iguaçu river is filled with sediments from the Serra do Mar, mainly by Quaternary alluvium deposited by the tributaries of this river. Fig 2 illustrates the perspective of the local relief, showing the Serra do Mar in the background, which contributes to the collection of water from the floodplains and the contribution of sediments to the plains.

**Fig 2.**
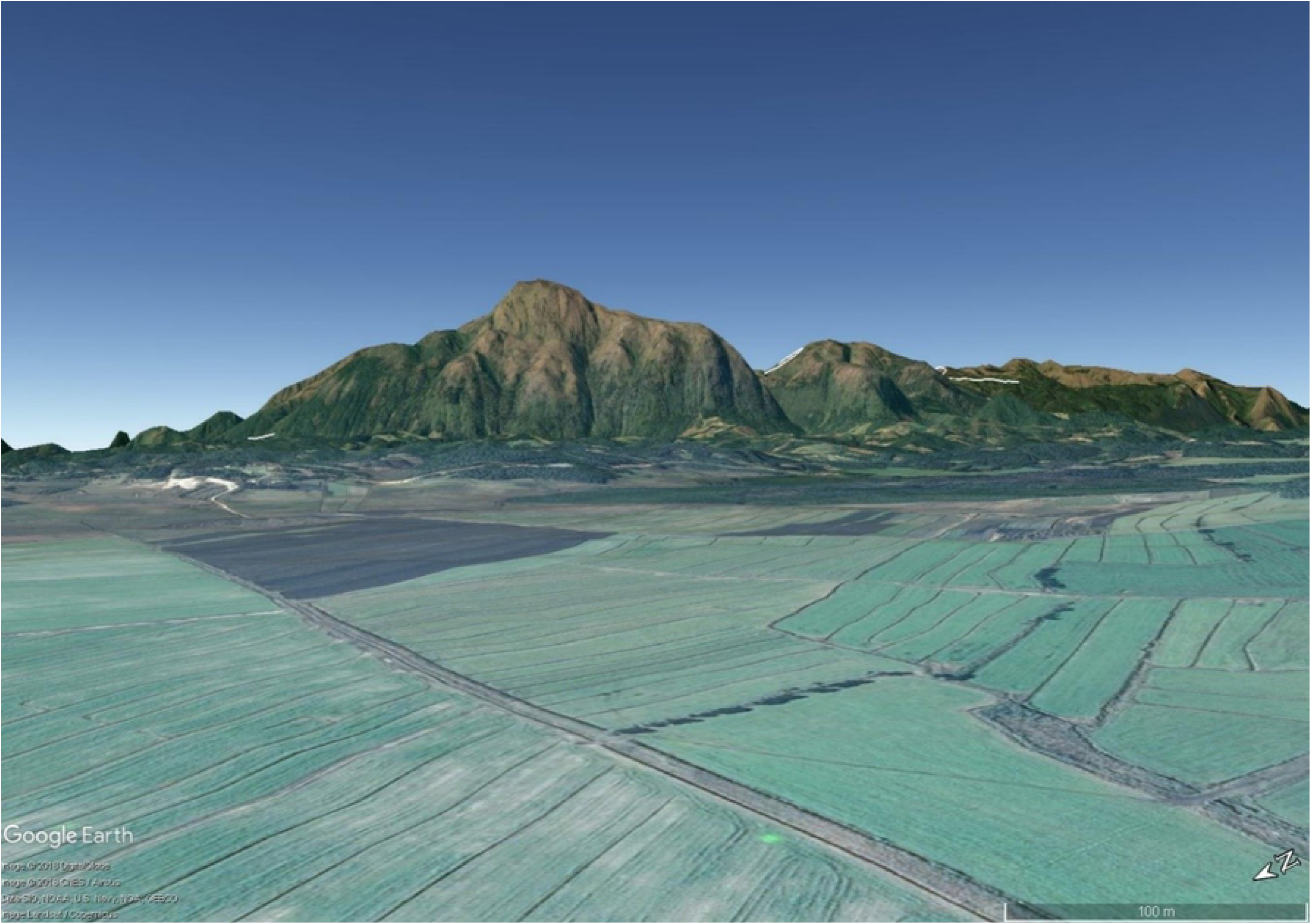
Satellite image in elevation perspective. The image illustrate the contribution of the Serra do Mar to the plain of the Upper Iguaçu river.

The local climate is classified as Cfb - humid subtropical mesothermal - according to Köppen [25]. The region is under the intense influence of relief rains (orographic) due to its proximity to the Serra do Mar, which guarantees a humid and cold climate for a long period of the year. The local vegetation is classified as Ombrophylous Mixed Forest, with the presence of Araucaria angustifolia and fields.

### Sampling

To relate the formation processes of kaolin deposits and geomorphological variations, samples were collected at two points. The first is situated in the plain of the upper Iguaçu river, in the municipality of Tijucas do Sul, coordinates 25°55′28.47″ S; 49°12′1.95″ W, at an altitude of 930 m above sea level. This area was chosen because it presents little anthropic influence and, thus, preserves the original characteristics of the sampled material (Fig 3).

**Fig 3.**
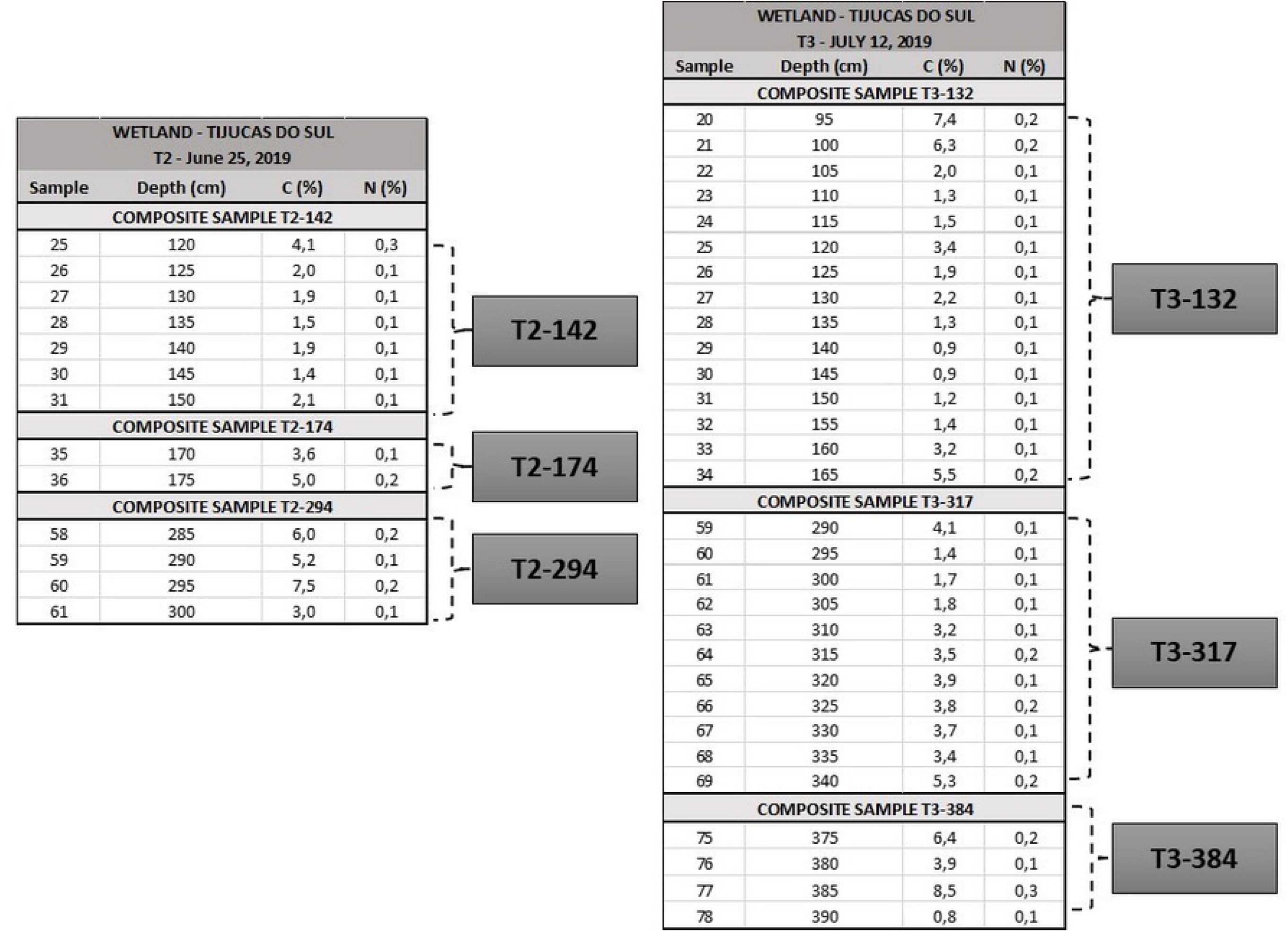
Sample separation scheme for mineralogical analysis

To define the point to be sampled, we surveyed the depth of the peat bog on the Upper Iguaçu plain, using a 6 m long tube and GPS/GNSS. Observations were made at approximately 10 m spacing, totaling 21 points sampled.

The peat volume was calculated with the *Posição* software. From this result, the collection area was estimated in the deepest region, in the center of the peat bog, sampling 2 sites of greater depths, 30 m apart from each other. At these sites, tubes were sampled using a 6 m long aluminum tube, inserted with the aid of a vibracore, to attenuate sample compaction [26]. Each core was opened and the sample inside the tube was sectioned every 5 cm. For comparison with possible kaolin from a sediment source area, two other points were sampled in an area of the “Serra do Caulim” mining company, geographic coordinates 25°34′53.67″ S; 48°59′00.71″ W and altitude 947 m from sea level. This area is situated amidst the hills of the Serra do Mar, with rocks of the gneiss-migmatite complex (Fig 1). At this site, one sample was collected in the mining area and another in the regolith, exposed by a ravine.

### Laboratorial Analysis

To better separate the mineral material from the organic material in the samples from the upper Iguaçu, the total C contents were determined via dry combustion with the Elementar Vario EL III equipment.

Samples with total C contents lower than 8%, associated with light coloration (white, gray, or light brown), were used for grain size and mineralogy analyses. A total of 43 mineral subsamples were selected, which formed 6 composite samples as shown in Fig 3, identified according to their collection tube and average depth. Thus, sample T2-142 refers to tube 2 and average depth of 142 cm, and so on (Fig 4).

**Fig 4.**
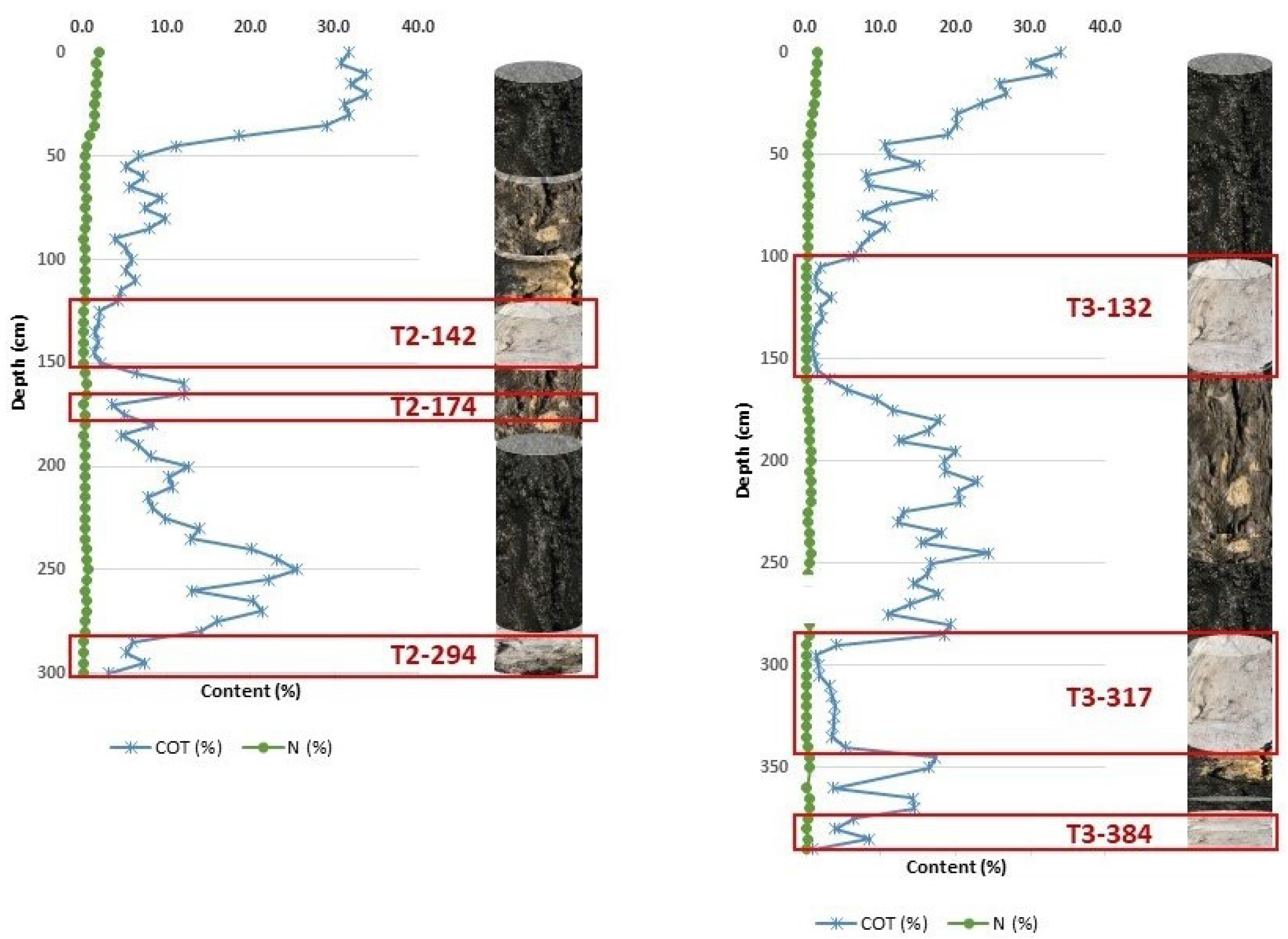
Total organic carbon (TOC) and N contents in the sampled.

The organic material from all samples was removed with a 30% (v/v) H_2_O_2_ solution in a water bath at about 60°C. Subsequently, particle size analysis was performed by the pipette method [27]. For the mineralogical analyses, the fine soil samples were dispersed with 0.5 mol L^−1^ NaOH solution, and the sand fraction was separated by sieving, and the clay was separated from the silt by siphoning, according to Stokes’ law.

### Mineralogical Analysis

The sand and silt fractions of all samples were obtained analyzed in the Panalytical X’Pert^3^ Powder model X-ray diffractometer, with a Cu Kα radiation source, with sweep from 3 to 65° 2θ, operating at 40 mA and 40 kV.

For the clay fraction, prior to X-ray analysis, extraction of the pedogenic Fe oxides with dithionite-citrate-bicarbonate (DCB) solution was performed in four successive extractions [28]. The extraction residues were washed and then frozen for freeze-drying. The mean crystal diameter (MCD) of kaolinite was calculated from the width at half height (WL) of reflections (001), (200) and (060), using the Scherrer equation [23, 29]:

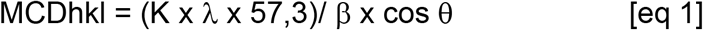

MCDhkl = DMC perpendicular to the plane hkl (nm);

K = 0,9 (constant);

λ = wavelength of the radiation used (0.15405 nm);

θ = Bragg’s angle;

57,3 = factor transforming the value of graus for radians;

β = corrected width at mid-height (WMH - °2θ).

To correct the corrected WMH of the peak of interest, Si standard was used to obtain the WMH due to instrumental distortion. To identify possible interstratification of kaolinite with 2:1 mineral, the asymmetry index (AI) was determined at the equivalent reflectance to the plane (001) of kaolinite [30]. The index of crystallinity (HBCI) of Kaolinite was also evaluated according to Hughes and Brown [24], with the equation:

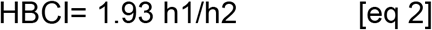

h1 is the peak height (020) of the kaolinite

h2 is the difference between triplet’s background (around 0.26 e 0.23 nm) and general background.

The Kt and Gb contents of all samples (plain and Serra do Mar) of iron-free clay (treated with DCB), were determined by thermogravimetric analysis (TDA/TG) in Shimadzu equipment and DTG-60 model, simultaneous ATG-TG APPARATUS. The samples were subjected to heating from room temperature to 950°C in a platinum crucible, at a heating rate of 10 °C min^−1^ and Nitrogen gas flow rate of 50 mL min^−1^.

Samples T2-294, T3-317, T3-384 and SAP, in the natural silt and clay fractions, were separated for analysis in the TESCAN VEGA3 LMU Scanning Electron Microscope with 3 nm resolution, with EDS type chemical analysis (Oxford) and AZ Tech software (Advanced). These samples had higher Ct peaks in the silt fractions, as well as 2:1 clay peak and were therefore subjected to microscopic analyses to complement the study.

The clay and silt fraction data were evaluated separately by means of Principal Component Analysis (PCA) with the software “R Studio” (RStudio, Boston, MA, USA). In order to verify the degree of similarity between the samples, cluster analysis by the minimum Euclidean distance was also performed.

## Results and Discussion

The boreholes in the plain indicated that the deepest depth of the peat bog was 4 m. With interpolation of the values, the volume of this bog was estimated at 38,514 m^3^ in an area of 21.066 m^2^, which could store up to 7,500 m^3^ of water in the first 50 cm depth [31]. At the base of the tube, at approximately 400 cm, the common presence of sand-sized grains was observed, with whitish or translucent coloration, suggesting the presence of feldspar or quartz.

Through nitrogen and total organic C (TOC) analysis, the regions of mineral material for mineralogical analysis (organic C < 8%) were estimated (Fig 4), concomitant with the coloration aspect, and three sections of the tubes were found with kaolinitic material deposited alternately with the organic material, suggesting that the deposition of kaolin in the region occurred in several pediplaning events.

In the cores collected in the plain of the upper Iguaçu river, the particle size analysis showed great variation between the proportions of the particles (Table 1), a typical behavior of deposition environments [32]. These variations in particle size composition between samples from the same point, coupled with clear transition between lighter and darker colored materials, are evidence of discontinuity, where the materials were deposited as packages in different events.

**Table 1.** Granulometry of saprolite samples (SAP), Kaolinitic deposit of serra do mar (CAU) and tubes from plain of the Upper Iguaçu river (T2 and T3).

| Sample | Sand (%) | Clay (%) | Silt (%) | Silt/Clay |
| --- | --- | --- | --- | --- |
| **SAP** | 20.6 | 22.1 | 57.3 | 2.60 |
| **CAU** | 30.4 | 20.8 | 48.8 | 2.34 |
| **T2-142** | 8.9 | 46.0 | 45.1 | 0.98 |
| **T2-174** | 6.3 | 36.4 | 57.3 | 1.58 |
| **T2-294** | 12.4 | 51.5 | 36.1 | 0.70 |
| **T3-132** | 10.1 | 49.4 | 40.5 | 0.82 |
| **T3-317** | 28.2 | 46.5 | 25.3 | 0.54 |
| **T3-384** | 4.4 | 47.2 | 48.4 | 1.03 |

### Thermo-differential Analysis and Thermo-gravimetric (TDA/TG)

The results obtained by means of TDA/TG (Table 2) show the predominance of kaolinite in the clay fraction of the samples. The highest kaolinite contents were found in core sample 3 (T3-132), with approximately 850 g kg^−1^, while the lowest contents were observed in the Serra do Mar saprolite (SAP), close to 550 g kg^−1^. Comparing the two samples, the saprolite has around 66% (2/3) of the kaolinite found in the floodplain areas. The Serra do Mar region, because it is affected by orographic rains, is submitted to an intense process of desilication and kaolinite formation [18], resulting in relatively high values of this mineral.

**Table 2.** Kaolinite (Kt) and Gibsite (Gb) content and respective dehidroxilation temperature.

| Sample | Gb<sup>1</sup> | T °C<sup>2</sup> | Kt<sup>1</sup> | T °C<sup>2</sup> |
| --- | --- | --- | --- | --- |
| SAP | 0.0 | - | 549 | 501 |
| CAU | 0.0 | - | 676 | 500 |
| T2-142 | 78 | 250 | 764 | 492 |
| T2-174 | 48 | 248 | 504 | 477 |
| T2-294 | 91 | 252 | 731 | 492 |
| T3-132 | 127 | 253 | 845 | 484 |
| T3-317 | 112 | 255 | 799 | 487 |
| T3-384 | 76 | 249 | 814 | 493 |

<sup>1</sup> Mineral content in the iron-free clay fraction. <sup>2</sup> dehydroxilation temperature.

Still according to Furian et al. [18], the environmental conditions of the Serra do Mar can result in the formation of significant amounts of Gibbsite. However, the results obtained with TDA/TG revealed the absence of this mineral in the Serra do Mar samples. In the lowlands (T2 and T3), Gibbsite occurs in small amounts, with a maximum of 127 g kg^−1^ (Table 2). Some of the Gibbsite found in the lowlands may be a result of the desilication process after sediment deposition in the wetter climate of the current conditions [35], as well as some of the kaolinite found in these areas.

The dehydroxylation temperature has a strong and positive correlation with the size and degree of crystallinity of minerals [23]. The temperatures at which the dehydroxylation peak of kaolinite occurred were higher for the samples from Serra do Mar (Table 2), suggesting larger crystals of this mineral, which may be related to the environment with less interference from other components, such as organic material.

Considering only the kaolinite contents in the samples of both areas, it was not possible to infer about the genesis of the minerals in the floodplain area, considering that in hydromorphic conditions there is the removal of Fe oxides. As a consequence of the removal of oxides, a residual concentration of kaolinite may have occurred, resulting in the higher values observed in the cores (T2 and T3). Moreover, the lower kaolinite contents in the SAP may be associated with the higher quartz contents in the clay fraction, observed by the peak at 3.44 nm (26.6° 2θ - Fig 5). The presence of quartz in the clay fraction is common for acidic rocks, such as granites and sandstones [33]. When in the clay fraction, quartz has high solubility, increasing the Si content in solution, which can slow down the desilication process in the saprolite. Moreover, part of the silica removed from the SAP can be concentrated in the adjacent area and form kaolinite, as occurs in the kaolinitic deposit (CAU) of Serra do Mar [34].

**Fig 5.**
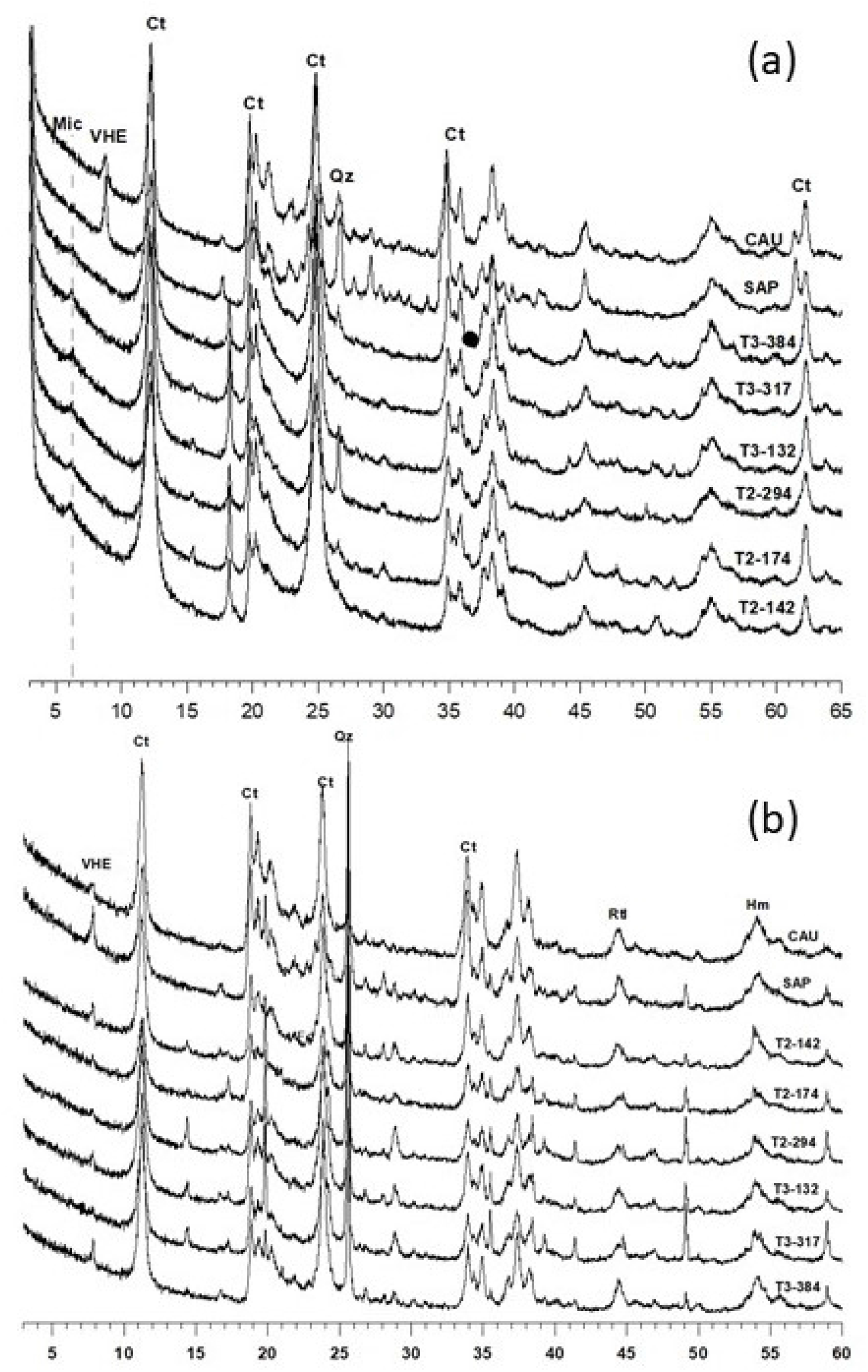
X-ray diffractogram for non-oriented samples after iron extraction by DCB: a) clay and b) silt fraction.

### X-ray diffraction

In the samples of the silt fraction there was a predominance of quartz, however, with significant participation of kaolinite (Fig 5). On the other hand, in the clay fraction was observed the predominance of kaolinite, with the presence of quartz, identified in the peak of 3.44 nm. The Gibbsite peaks in the XRD patterns of the cores were consistent with the thermal analysis data (Table 2). The peak at 1.0 nm shows significant presence of mica in the Serra do Mar samples (Fig 5). Mica was also observed in all samples of the silt fraction, both in the seamount and core samples.

### MEV-EDS da fração silte

For SEM-EDS analysis, samples of the saprolite (SAP), the kaolin interlayer of core 2 and the two kaolin layers found at greater depth in core 3 were selected. The silt fraction was analyzed to investigate the origin of the kaolinite identified in the XRD.

In almost the entire area of the SEM images it is possible to observe crystals with planar growth, common in phyllosilicate minerals (Fig 6). Some structures have larger openings between the layers, which facilitates the movement of water through the material. From the results of the EDS microanalysis (Table 3), it can be seen that the Si/Al ratio for most of the samples is in the range of 1.0 to 1.2, a common ratio for 1: 1 clay minerals. The K values were higher in the Saprolite samples, but similar values were also found at some points in the samples of core core sample 3. The images associated with the values obtained by EDS suggest that the structures observed are pseudomorphs, where kaolinite uses the structure of micas and forms the stacking of crystals by diagenesis, occurring simultaneously the removal of K [9, 14, 16, 21].

**Fig 6.**
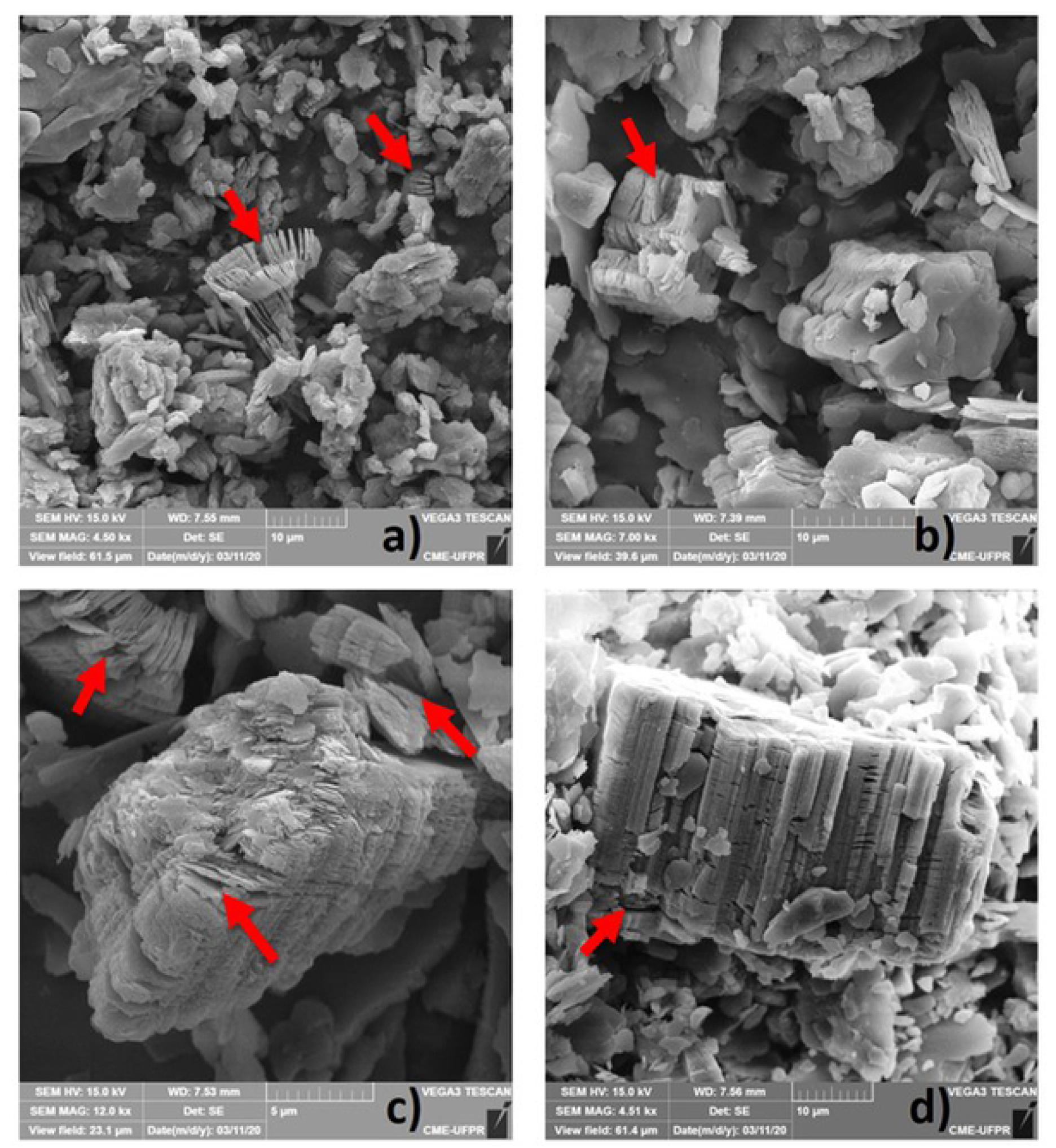
Micrographs of the silt fractions and the pseudomorphs. a) Saprolite (SAP); b) T2-294; c) T3-317 and d) T3-384. The arrows in red indicate the transformations from muscovite into kaolinite.

**Table 3.** Semi-quantitative contents of the elements assessed through energy dispersive spectroscopy (EDS).

|  | O | Na | Mg | Al | Si | K | Ca | Ti | Fe | Cu | Si/Al |
| --- | --- | --- | --- | --- | --- | --- | --- | --- | --- | --- | --- |
| SAP | 47.95 | 0.11 | 0.33 | 16.32 | 23.8 | 3.57 | - | 0.14 | 1.48 | 0 | 1.46 |
|  | 48.74 | 0.04 | 0.53 | 11.23 | 17.72 | 3.36 | 0.03 | 8.6 | 1.83 | 0.1 | 1.58 |
|  | 44.66 | 0.13 | 0.23 | 14.57 | 21.8 | 3.38 | 0.05 | 4.45 | 1.96 | 0 | 1.50 |
|  | 47.93 | 0.12 | 0.29 | 13.21 | 19.53 | 2.82 | 0.05 | 5.99 | 1.67 | 0.13 | 1.48 |
|  | 48.66 | 0.1 | 0.25 | 9.22 | 31.2 | 2.13 | 0.03 | - | 0.79 | 0.01 | 3.38 |
| T2-294 | 46.85 | 0.13 | 0.15 | 14.82 | 18.37 | 1 | 0 | 1.31 | 3.58 | 0.01 | 1.24 |
|  | 47.53 | 0.14 | 0.14 | 15.01 | 17.99 | 1.14 | 0 | 1.25 | 3.23 | 0 | 1.20 |
| T3-317 | 51.29 | 0.2 | 0 | 16.21 | 15.94 | 0.69 | 0.12 | 1.03 | 2.88 | 0 | 0.98 |
|  | 52.22 | 0.15 | 0.15 | 19.47 | 22.92 | 1.37 | 0.07 | 1.27 | 2.07 | 0 | 1.18 |
|  | 51.88 | - | - | 13.71 | 25.53 | 0.91 | - | 1.05 | 1.79 | 0 | 1.86 |
|  | 51.13 | 0.18 | 0.19 | 19.73 | 23.57 | 0.68 | - | 1.06 | 3.12 | 0 | 1.19 |
|  | 50 | 0.2 | - | 17.81 | 21.86 | 2.56 | - | 1.09 | 2.59 | 0.1 | 1.23 |
|  | 49.55 | 0.24 | 0.11 | 18.2 | 19.92 | 0.68 | 0.1 | 1.36 | 2.61 | 0 | 1.09 |
|  | 47.97 | 0.17 | 0.13 | 15.61 | 14.73 | 0.72 | 0.17 | 1.28 | 3.69 | 0 | 0.94 |
|  | 50.64 | 0.15 | 0.18 | 17.65 | 21.85 | 0.79 | 0.04 | 1.33 | 2.49 | 0 | 1.24 |
|  | 46.88 | 0.18 | 0.52 | 17.37 | 22.13 | 2.99 | 0 | 1.09 | 4.03 | 0 | 1.27 |
| **T3-384** | 52.26 | 0.07 | 0.15 | 18.15 | 18.99 | 0.83 | 0 | 0.7 | 1.62 | 0.01 | 1.05 |
|  | 46.06 | 0.06 | 0.2 | 18.6 | 19.47 | 1.54 | 0 | 2.92 | 3.86 | 0 | 1.05 |
|  | 45.07 | 0.13 | 0.14 | 19.73 | 20.75 | 2.64 | 0 | 1.29 | 2.47 | 0 | 1.05 |
|  | 45.69 | 0.12 | 0.13 | 12.11 | 12.24 | 0.96 | 0 | 22.55 | 2.21 | 0.03 | 1.01 |
|  | 47.17 | 0.13 | 0.09 | 16.54 | 12.72 | 1.04 | 0.29 | - | 3.2 | 0 | 0.77 |
|  | 51.57 | 0.21 | 0.14 | 17.28 | 18.57 | 1.08 | 0 | 2.43 | 2.18 | 0.04 | 1.07 |
|  | 49.59 | 0.12 | 0.18 | 19.91 | 21.38 | 1.18 | 0 | 0.48 | 1.45 | 0 | 1.07 |
|  | 50.6 | 0.15 | 0.13 | 19.35 | 21.15 | 1.17 | 0 | 0.53 | 1.59 | 0.12 | 1.09 |
|  | 49.24 | 0.14 | 0.51 | 18.23 | 20.3 | 2.94 | 0.11 | 0.55 | 2.53 | 0 | 1.11 |
|  | 50.58 | 0.05 | 0.3 | 19.31 | 20.43 | 1.07 | 0.05 | 0.58 | 1.46 | 0 | 1.06 |

The presence of pseudomorphic Mc-Kt in the silt fraction of all analyzed samples makes it difficult to define the genesis of the floodplain sedimentary packages. The three genesis premises admit the presence of pseudomorphic Mc-Kt in both the Serra do Mar and the plain: i) the pseudomorphic Kt was formed in the Serra do Mar and transported to the plain. In this situation the pedobioturbation conditions were not intense enough to break the pseudomorphs and carry the Kt particles to the clay fraction; ii) the pseudomorphic mica kaolinite in the floodplain area is autochthonous, formed directly from the weathering of mica transported to the floodplain area; iii) a mixed situation, where part of the pseudomorphic kaolinite was transported and part was formed in situ in the floodplain, by mica weathering.

In this sense of fragmentation of the pseudomorphs as a function of transport, it can be inferred that the structures shown in Fig 6, especially in sample 3, were not transported from the highest part of the Serra do Mar. The thickness of the pseudomorph highlighted in sample T3-384 is about 30 μm (stacking of kaolinite particles in basal direction) and the highlighted particle in the SAP is about half that thickness. As in the transport process there is only the possibility of breaking the pseudomorphs, it can be inferred that the kaolinite with this morphology in the silt fraction of core sample 3 was formed in situ by the weathering of the transported mica. On the other hand, the thickness of the pseudomorph kaolinite is more compatible with the SAP.

The results obtained by microchemical analysis (EDS) of the kaolinite crystals in the silt fraction also allow us to infer about its genesis in the floodplain areas (Table 3). The highest values of are related to residual K in the mica, being a strong indication of the presence of residual mica layer in the pseudomorphic kaolinite particles. The higher K content in the SAP sample cannot be considered as a conclusive result, because the K in these samples may be removed during the process of transport and deposition of kaolinite in the floodplain. The difference in Fe content between the SAP and the cores reinforces that the kaolinite in the floodplain silt was formed in situ by weathering of the mica. The higher Fe content of kaolinite in the silt fraction of the floodplain is also evidence of in situ formation of this mineral in the floodplain. Possibly, there was a greater contribution of biotite particles in the formation of the silt-sized Kt in the plain. If silt was transported from higher positions, the iron content in the Kt should be lower, similar to what was observed for K, and not higher in the floodplain samples.

Consolidated with the microscopy data, it is possible to infer that the silt-sized Kt in the plain soils were formed in situ by the diagenesis of Mc particles (biotite participation). Possibly, these silt-sized mica particles were transported and deposited in the floodplains.

### Kaolinite cristalography

With the exception of the Serra do Mar saprolite clay fraction, the samples that had the highest values for the HBCI, had the lowest calculated values for the Asymmetry Index (AI) of the (001) plane of Kt (Table 5). There is a direct relationship between AI and interstratification with 2:1 layers in Kt. One of the factors that reduces the crystallinity of Kt is the intensity of these interstratifications [36]. The HBCI values of the samples from tube 3 (T3) were similar to those observed for the samples from the Serra do Mar, while the samples from tube 2 (T2), showed lower values. The kaolinite from the SAP presents more interstratifications with 2:1 mineral (AI = 0.213) than the kaolinite from the CAU (AI = 0.075). This behavior indicates that the genesis of the kaolinite of the clay fraction in the saprolite and the kaolin mine in the Serra do Mar may be partially different.

**Table 5.** Mean diameter of the crystal (MCD). Number of plans and dimension relationships between the plans of the kaolinite crystals for the silt and clay fractions.

|  | Silt |  |  |  |  |  | Clay |  |  |  |  |
| --- | --- | --- | --- | --- | --- | --- | --- | --- | --- | --- | --- |
|  | plan | MCD (nm) | #plan | a/c<sup>1</sup> | b/c<sup>2</sup> | a/b<sup>3</sup> | MCD (nm) | #plan | a/c | b/c | a/b |
| CAU | (001) | 9.5 | 13 | 1.8 | 1.8 | 1.0 | 11.0 | 15 | 1.1 | 1.6 | 0.7 |
|  | (200) | 17.0 | 68 |  |  |  | 12.5 | 50 |  |  |  |
|  | (060) | 16.9 | 113 |  |  |  | 17.6 | 118 |  |  |  |
| SAP | (001) | 6.1 | 9 | 1.9 | 2.1 | 0.9 | 10.0 | 14 | 1.3 | 1.8 | 0.7 |
|  | (200) | 11.5 | 46 |  |  |  | 12.7 | 51 |  |  |  |
|  | (060) | 12.8 | 86 |  |  |  | 18.2 | 122 |  |  |  |
| T2-142 | (001) | 6.1 | 9 | 2.0 | 2.9 | 0.7 | 8.2 | 11 | 2.0 | 1.8 | 1.2 |
|  | (200) | 12.5 | 50 |  |  |  | 16.8 | 67 |  |  |  |
|  | (060) | 17.8 | 120 |  |  |  | 14.5 | 97 |  |  |  |
| T2-174 | (001) | 6.9 | 9 | 2.2 | 1.7 | 1.3 | 7.8 | 11 | 1.8 | 1.8 | 1.0 |
|  | (200) | 15.2 | 61 |  |  |  | 13.7 | 55 |  |  |  |
|  | (060) | 11.4 | 77 |  |  |  | 13.9 | 93 |  |  |  |
| T2-294 | (001) | 7.5 | 10 | 1.6 | 2.4 | 0.7 | 9.3 | 13 | 2.1 | 1.8 | 1.2 |
|  | (200) | 12.0 | 48 |  |  |  | 19.8 | 79 |  |  |  |
|  | (060) | 17.8 | 120 |  |  |  | 16.9 | 113 |  |  |  |
| T3-132 | (001) | 6.0 | 8 | 1.9 | 2.7 | 0.7 | 9.0 | 13 | 1.6 | 2.0 | 0.8 |
|  | (200) | 11.5 | 46 |  |  |  | 14.4 | 58 |  |  |  |
|  | (060) | 16.0 | 107 |  |  |  | 17.9 | 120 |  |  |  |
| T3-317 | (001) | 6.4 | 9 | 1.8 | 2.6 | 0.7 | 9.3 | 13 | 1.7 | 1.8 | 1.0 |
|  | (200) | 11.5 | 46 |  |  |  | 16.1 | 65 |  |  |  |
|  | (060) | 16.9 | 113 |  |  |  | 16.9 | 114 |  |  |  |
| T3-384 | (001) | 7.7 | 11 | 2.1 | 2.3 | 0.9 | 13.1 | 18 | 1.3 | 1.4 | 1.0 |
|  | (200) | 16.0 | 64 |  |  |  | 17.2 | 69 |  |  |  |
|  | (060) | 17.8 | 120 |  |  |  | 17.9 | 120 |  |  |  |

<sup>1. 2. 3</sup> The relations were calculated from the mean crystal diameter (MCD) for the planes (200) - dimension a; plane (060) - dimension b; and plane (001) - dimension c. The number of plans was calculated by dividing the DMC by the distance between plans (d).

The intense presence of pseudomorphic kaolinite in the silt fraction in the SAP (FIGURE 19) it can be inferred that the kaolinite in the clay fraction of the SAP was also partly formed by diagenesis of mica particles. This process preserves layers of mica in the kaolinite structure. As for kaolinite from the Serra do Mar (CAU) kaolin mines, it is more accepted that it was formed and concentrated from the neogenesis of potassic feldspar veins present in the migmatites [37], which reduces the AI of the mineral.

The higher microstrain values (measure of structural deformation) for the (060) plane of kaolinite, may be associated with the increased isomorphic substitution of Al^3+^ by Fe^3+^ in the octahedral lamina [38]. In the present work, the highest values were observed for the samples of tube 2 and, in the case of T2-174, they reached 0.52 (TABLE 5). In the clay fraction, the entry of Fe into the octahedral lamina (increased microstrain) was evidenced by the increase in the unit cell size (R^2^ between microstrain (200) and d(200) = 0.61) (Fig 7). Fe^3+^ has a larger ionic radius than Al^3+^ and the isomorphic substitution of these elements in the octahedral lamina promotes an increase in the unit cell size of kaolinite [39]. Despite the increase in unit cell dimensions in the x- and y-plane (unit cell dimensions a and b), the entry of Fe^3+^ into the octahedral lamina promoted a reduction in kaolinite thickness, showing a negative relationship between microstrain (002) and d(002) (Fig 7). The repulsion between the trivalent cations of two consecutive octahedral positions promotes distension and shortening of the shared edge in the octahedral lamina, and as a result there is a corrugation and reduction of the thickness of this lamina in kaolinite [22].

**Fig 7.**
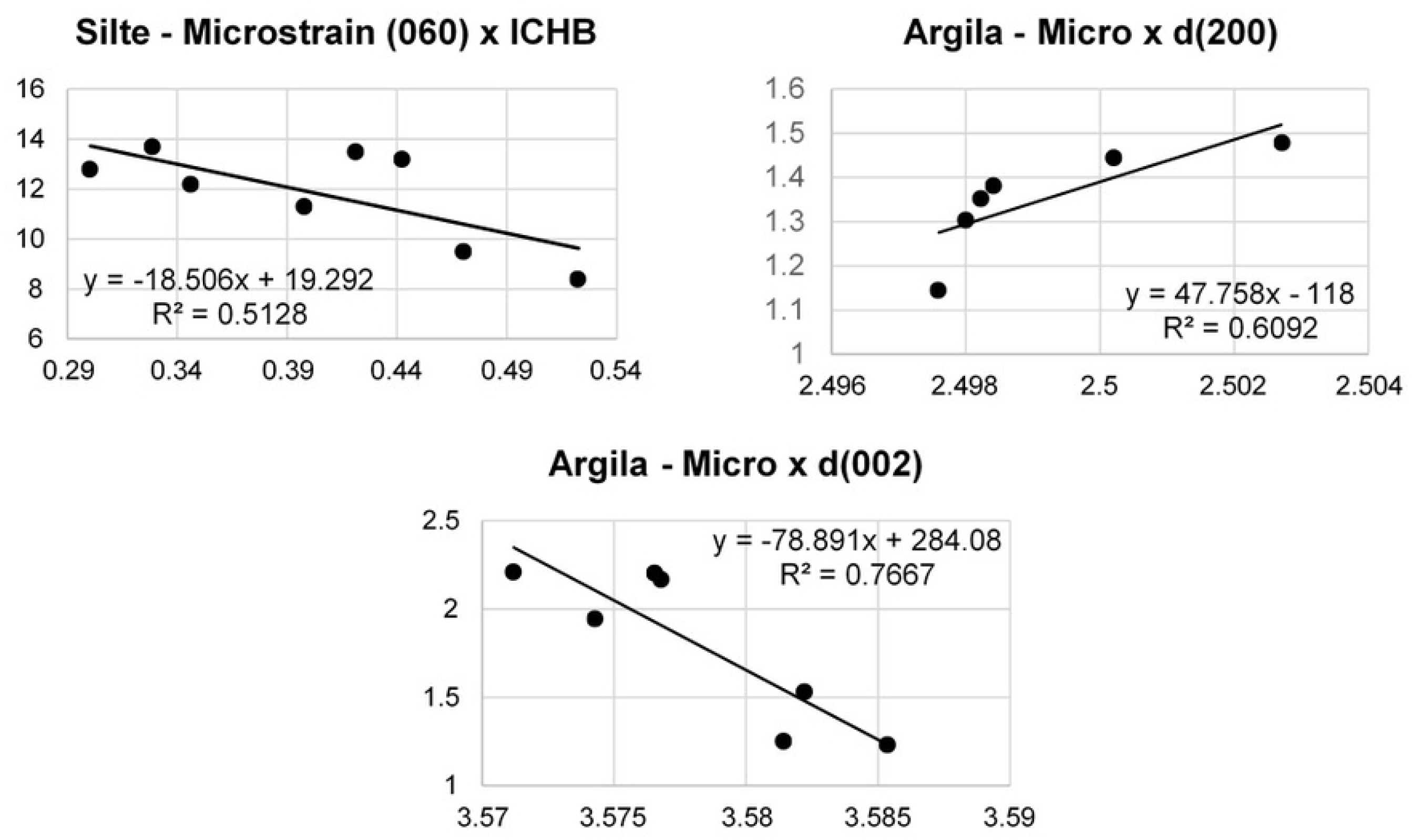
Relationships between the crystallographic attributes of kaolinite.

Considering the mean crystal diameter (MCD), the clay samples presented higher values than the silt fraction crystals (Table 5). This indicates that for the kaolinite particles to reach the silt size an agglomeration of particles was necessary, in the form of a pseudomorph structure of mica or cemented and stable aggregates [23]. The relations between the diameters of the crystals in the three-dimensional axes demonstrate the preferential growth in planes a and b at approximately twice the growth in dimension c. This pattern is typical of phyllosilicate minerals [22], and were observed both in the crystals of the silt fraction and the clay fraction. Considering this attribute alone, there is no way to differentiate kaolinite from the different samples collected.

### Statistical Analysis

In the PCA for the attributes of the clay fraction, it is observed that the first axis opposes the contents of gibbsite and the HBCI of kaolinite, indicating that the weathering process is the main responsible for the variability of these data, accounting for 40.9% of all variation (Fig 8). Factor axis 2, on the other hand, explains 23.84% of the variation in the data, opposing the AI of kaolinite to the DMC of the plane (001). In Fig 8 the contribution of the samples to the factorial axes are represented, demonstrating that the samples from the Kaolin Mountains (SAP and CAU) are positioned on the same side as the HBCI. This behavior is related to the lower degree of weathering of the material collected from the Kaolin Mountains.

**Fig 8.**
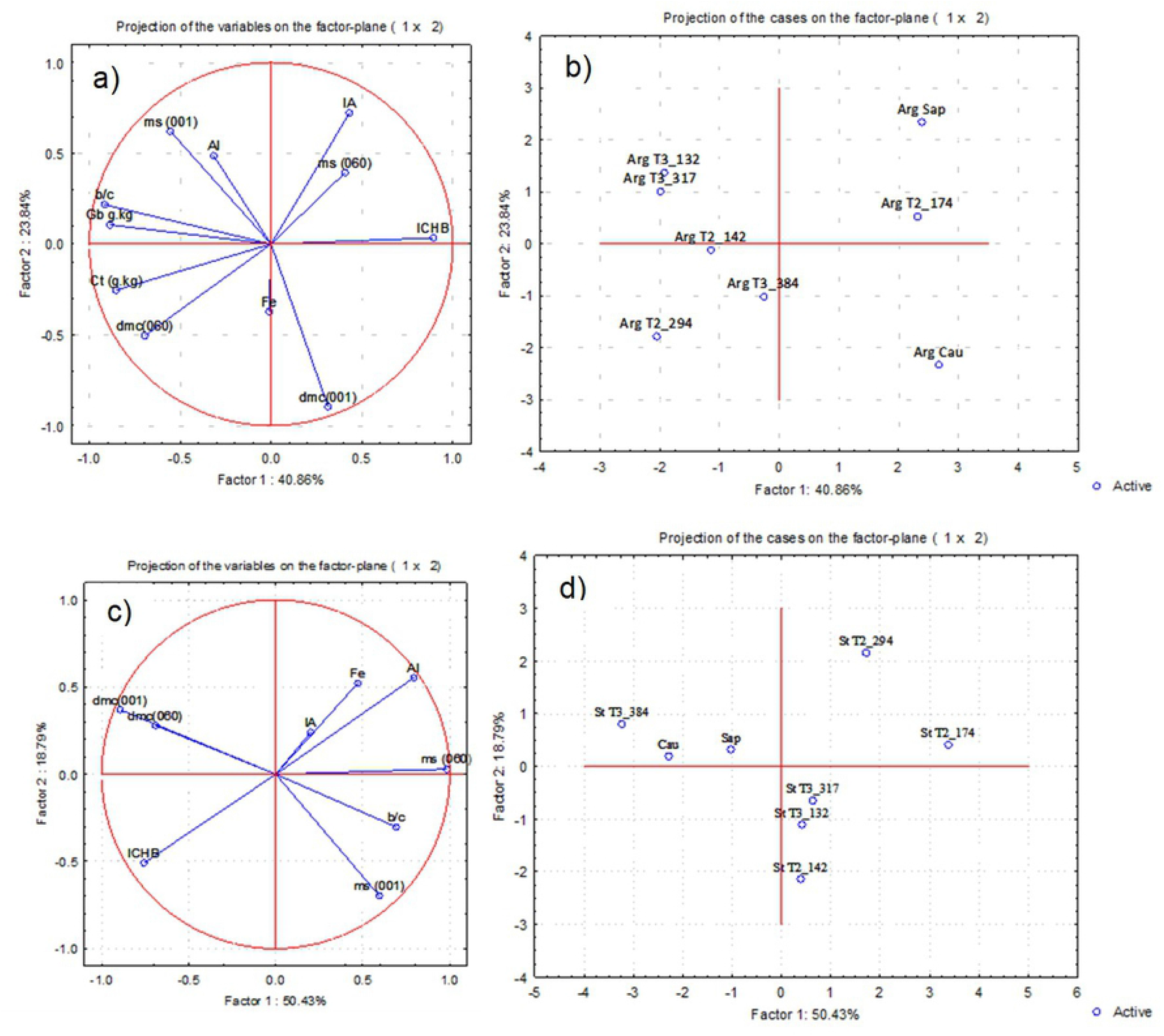
Principal component analysis of the clay (a, b) and silt (c, d) fractions.

The first PCA factorial axis for the silt fraction of the samples contemplates 50.4% of the total variability of the data and opposes the HBCI to the Al, Fe, kaolinite AI and microstrain contents in the (060) kaolinite plane (Fig 8), while the second factorial axis contemplates 18.8% of the variability of the data and opposes the HBCI and the Fe, Al contents and the kaolinite AI. The projection of the samples on the factorial plane reinforces the processes discussed for the clay fraction. The opposition of the HBCI to the Al, Fe and AI on axis 2 suggests that in addition to the weathering process represented from axis 1, there is an increase of Fe in the mineral, resulting from the isomorphic substitution of Al^3+^ by Fe^3+^, also reflecting in the microstrain values for the (060) plane.

As a way to compare between the samples and establish the similarities between them, cluster analysis was performed [38]. In the results of this analysis, it was observed that the samples from the Serra do Mar have high similarity, differentiating themselves from the others at a higher level (Fig 9). However, the samples from the Serra do Mar are most similar to the deepest sample from tube 3 (T3-387). This similarity may be a result of the lesser influence of pedogenetic processes (and organic material), or may represent the greater similarity with the parent material, since this layer is closer to the basin base and bedrock.

**Fig 9.**
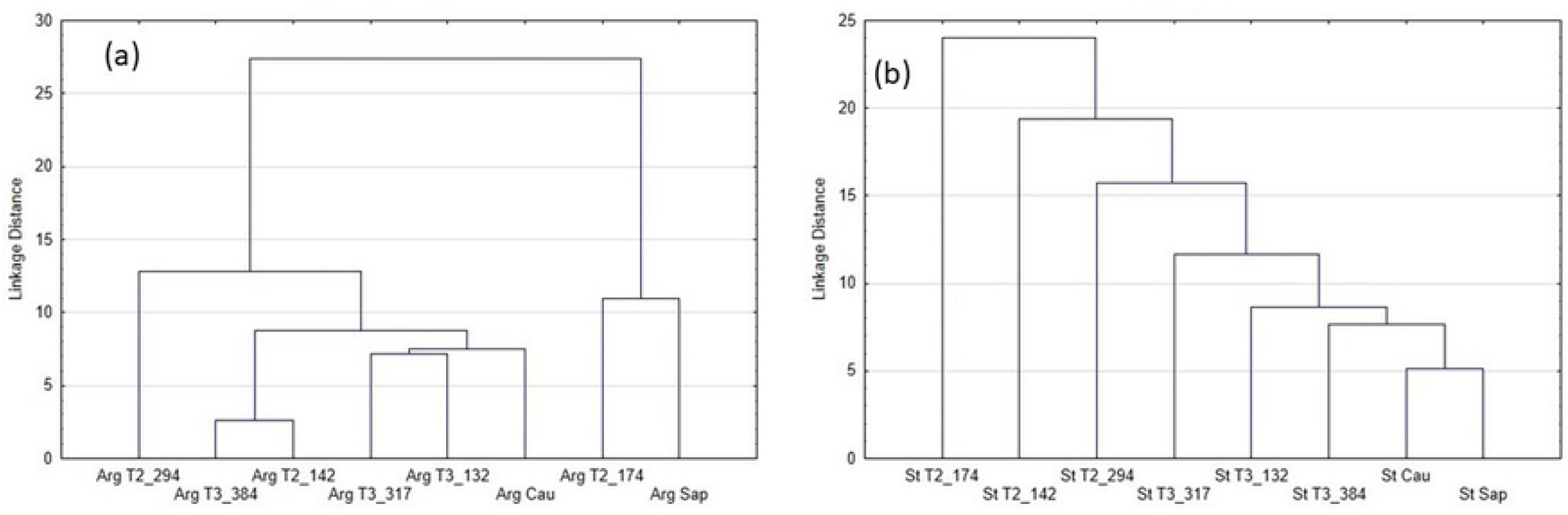
Cluster analysis of the Serra do Mar and plain of the Upper Iguaçu river.

For the clay fraction, the samples from the Serra do Mar were more distant from each other, being closer to other samples from the plain of the upper Iguaçu river (Fig 9). These results suggest that even some of the kaolinite present in the deposits is inherited from in situ weathering of the parent material [1]. This material, after passing through the processes of erosion, transport and deposition on the plain of the upper Iguaçu river, undergoes other weathering processes, but still preserving characteristics of its origin.

## Conclusions

The results obtained in this work indicate that the kaolinitic material found in the plains of upper Iguaçu river is the result of weathering processes in the floodplain itself, evidenced by the large pseudomorphs found, even larger than those observed in the Serra do Mar. Most of the kaolinite minerals present in this deposit were formed by the chemical weathering of the rocks of the crystalline basement, among them granites, gneisses and migmatites. However, part of them could have been formed in the Serra do Mar region and then transported to the lower part of the landscape by geomorphological processes of landscape flattening. Alternated to these geomorphological processes, pedological processes of paludization also occurred, resulting in the layers with high organic C content.

The participation of organic matter in soil acidification and kaolinite formation can be considered small, without enough expression to form the deposits found on the First Plateau of Parana, specifically on the upper Iguaçu plateau, especially in well drained areas. However, the elucidation of the geochemical and geomorphological events, in chronological and hierarchical order, are of great importance to establish the genesis model of the kaolinitic deposits.

## Aknowledgments

We thank CAPES (Coordination for the Improvement of Higher Education Personnel) for granting a scholarship (Master’s Degree) to conduct this study.

## References

1. Chen PY, Lin ML, Zheng Z. On the origin of the name kaolin and the kaolin deposits of the Kauling and Dazhou áreas, Kiangsi, China. Elsevier: Applied Clay Science, 1997.

2. Silva SP. Caulim. In: BRASIL, DEPARTAMENTO NACIONAL DE PRODUÇÃO MINERAL. Balanço Mineral Brasileiro 2001. Brasíla: DNPM, 2001. p.121–133.

3. Campos AP, Monteiro CC, Silva JPA, Silva RD. Caulim. In: Brasil, Departamento Nacional De Produção Mineral. Sumário Mineral Brasileiro. 2018. Available at: <http://www.anm.gov.br/dnpm/publicacoes/serie-estatisticas-e-economia-mineral/sumario-mineral/pasta-sumario-brasileiro-mineral-2018/caulim_sm_2018>. Acessed on mach 09th, 2020.

4. Mártires RAC. Caulim. In: BRASIL, Departamento Nacional De Produção Mineral. Economia Mineral do Brasil. DNPM. 2009; p.442–451.

5. Hinckley DN. Variability in “crystallinity” values among the kaolinites deposits of the coastal plain of Georgia and South Carolina. Clays and Clay Minerals. 1963; 11: 229–35.

6. Pruett R. Kaolin deposits and their uses: Northern Brazil and Georgia, USA. Applied Clay Science. 2016; 131:3–13. doi: 10.1016/j.clay.2016.01.048.

7. Sayin S. Origin of kaolin deposits: Evidence from the Hisarcik (Emet-Kütahya) deposits, western Turkey. Turkish Journal of Earth Sciences. 2007; 77–96.

8. Ismail S, Husain V, Anjum S. Mineralogy and Genesis of Nagar Parker Kaolin Deposits, Tharparkar District, Sindh, Pakistan. International Journal of Economic and Environmental Geology. 2014; 33–40. doi: 10.46660/ijeeg.vol11.iss1.1919.

9. Ekosse GIE. Kaolin deposits and occurrences in Africa: Geology, mineralogy and utilization. Applied Clay Science. 2010; 50:212–236. doi: 10.1016/j.clay.2010.08.003.

10. Gilg H, Weber B, Kasbohm J, Frei R. Isotope geochemistry and origin of illite-smectite and kaolinite from the Seilitz and Kemmlitz kaolin deposits, Saxony, Germany. Clay Minerals. 2003; 38:95–112. doi: 10.1180/0009855033810081.

11. Dill H, Bosse H, Henning K. Mineralogical and chemical variations in hypogene and supergene kaolin deposits in a mobile fold belt the Central Andes of northwestern Peru. Mineralium Deposita. 1997; 32:149–163. doi: 10.1007/s001260050081.

12. Montes CR, Melfi AJ, Carvalho A. Genesis, mineralogy and geochemistry of kaolin deposits of the Jari river, Amapá State, Brazil. Clays and Clay Minerals. 2002; 50:494–503. doi: 10.1346/00098600232051421

13. Wilson SW, Santos HP. Kaolin and halloysite deposits of Brazil. Clays and Clay Minerals. 2006; 41:697–716. doi: 10.1180/0009855064130213.

14. Galán E, Ferrel RE. Chapter 3 - Genesis os Caly Minerals. Developments in Clay Science. 2013; 5:83–126. doi: 10.1016/B978-0-08-098258-8.00003-1.

15. Murray H. Applied clay mineralogy today and tomorrow. Clay Minerals. 1999; 34: 39–49. doi: 10.1180/000985599546055.

16. Kotschoubey B. Cobertura Bauxítica e a Origem do caulim do Morro do Felipe, Baixo Rio Jari, Estado do Amapá. Revista Brasileira de Geociências. 1999; 29 (3): 331–338. doi

17. Biondi JC, Santos ER. Depósito de caulim de Tijucas do Sul (Mina Fazendinha, Tijucas do Sul - PR). Revista Brasileira de Geociências. 2004;. 34:243–252.

18. Furian S, Barbiero L, Boulet R, Curmi P, Grimaldi M, Grimaldi C. Distribution and Dynamic of Gibbsite and Kaolinite in an Oxisol of Serra do Mar, southeastern Brazil. Geoderma. 2002; 106:83–100. doi: 10.1016/S0016-7061(01)00117-3.

19. Mucha NM. Relação solo-relevo entre a Serra do Mar e Planalto do Alto Iguaçu como subsídio para o Mapeamento Digital de Solos. 139 f. Master Dissertation. Federal University of Parana, 2020.

20. Muggler CC. Polygenetic Oxisols on tertiary surfaces, Minas Gerais, Brazil-Soil genesis and landscape development. 1998. PhD Thesis, Wageningen Agricultural University, Wageningen, The Netherlands, 185p.

21. Kampf N, Curi N, Marques JJ. Intemperismo e Ocorrência de minerais no ambiente do solo. In: Melo VF, Alleoni LRF (eds.). Química e Mineralogia do Solo: Conceitos básicos. 2009; p. 333–379.

22. Melo VF, Wypych F. Caulinita e haloisita. In: Melo VF, Alleoni LRF (eds). Química e mineralogia do solo. Sociedade Brasileira de Ciência do Solo. 2009; p.427–504.

23. Melo VF, Singh B, Schaefer CEGR, Novais RF, Fontes MPF. Chemical and mineralogical properties of kaolinite-rich Brazilian soils. Soil Science Society of America Journal. 2001. 65:1324–1333. doi: 10.2136/sssaj2001.6541324x.

24. Hughes JC, Brown G. A cristallinity index for soil kaolinite and its relation to parent rock, climate and soil maturity. Journal Soil Science. 1979; 30:557–563.

25. Köppen W. Climatologia. México: Fundo de Cultura Econômica, 1931.

26. Martin L, Flexor JM, Suguio K. Vibrotestemunhador leve: construção, utilização e potencialidades. Revista do Instituto Geológico. 1995; 16:59–66. doi: 10.5935/0100-929X.19950004.

27. EMBRAPA. Análise Granulométrica – Dispersão Total. In: Manual de Métodos de Análise de Solo. Rio de Janeiro, Embrapa Solos. 2011. 2^a^ Ed. p. 43–49.

28. Mehra OP. Iron oxide removal from soils and clay by a dithionite-citrate system buffered with sodium bicarbonate. Minerals. 1960; 317–327.

29. Klug HP, Alexander LE. X-ray Diffraction Procedures for Polycrystalline and Amorphous Materials. John Wiley & Sons, New York (1954).

30. B. Singh, R.J. Gilkes. Concentration of iron oxides from soils clays by 5 M NaOH treatment: the complete removal of sodalite and kaolin. Clay Miner. 1991; 26: 463–472. doi: 10.1180/claymin.1991.026.4.02.

31. Cipriano-Silva R, Valladares GD, Pereira MG, Anjos LH. Caracterização de Organossolos em ambientes de várzea do nordeste do Brasil. Revista Brasileira de Ciência Do Solo. 2014; 38:26–38.dDoi: 10.1590/S0100-06832014000100003.

32. Lima F, Fernandes L, Melo M, Góes A, Machado D. Faciologia e contexto deposicional da Formação Guabirotuba, Bacia de Curitiba (PR). Brazilian Journal of Geology. 2013; 43: 168–184. doi: 10.5327/Z2317-48892013000100014.

33. Melo VF, Oliveira Junior JC, Batista AH, Cherobim VF, Favaretto N. Goethite and hematite in bichromic soil profiles of southern Brazil: Xanthization or yellowing process. Catena. 2020; 188, e: 104445. doi 10.1016/j.catena.2019.104445.

34. Singer A. The mineralogy of the clay fraction from basaltic soils in the galilee, Israel. Journal of Soil Science. 1966; 17:136–147. doi: 10.1111/j.1365-2389.1966.tb01461.x.

35. Chiapini M, Schellekens J, Calegari M, Almeida J, Buurman P, Camargo P, Vidal-Torrado P. Formation of black carbon rich ‘sombric’ horizons in the subsoil - A case study from subtropical Brazil. Geoderma. 2018; 314: 232–244. doi: 10.1016/j.geoderma.2017.10.031.

36. Aparicio P, Galán E. Mineralogical Interference on Kaolinite Crystallinity Index Measurements. Clays Clay Miner. 1999; 47:12–27. doi: 10.1346/CCMN.1999.0470102.

37. Oliveira MTG, Furtado SMA, Formoso MLL, Eggleton RA, Dani N. Coexistence of halloysite and kaolinite: a study on the genesis of kaolin clays of Campo Alegre Basin, Santa Catarina State, Brazil. Anais da Academia Brasileira Ciências. 2007; 79:665–681. doi: 10.1590/S0001-37652007000400008.

38. Prandel LV, Melo VF, Brinatti AM, Saab SC, Salvador FAS. X-ray Diffraction and Rietveld Refinement in Deferrified Clays for Forensic Science. Journal of Forensic Sciences. 2017, 62:251–257. doi: 10.1111/1556-4029.13476.

39. Awad ME, López-Galindo A, Sánchez-Espejo R, Sainz-Díaz CI, El-Rahmany MM, Viseras C. Crystallite size as a function of kaolinite structural order-disorder and kaolin chemical variability: Sedimentological implication. Applied Clay Science. 2018; 162:261–267. doi: 10.1016/j.clay.2018.06.027.

